# Pericyte-mediated constriction of renal capillaries evokes no-reflow and kidney injury following ischemia

**DOI:** 10.1101/2021.09.24.461675

**Authors:** Felipe Freitas, David Attwell

**Author notes:** **Address correspondence to:** David Attwell, Department of Neuroscience, Physiology & Pharmacology, Andrew Huxley Building, University College London, Gower Street, London, WC1E 6BT, UK.

## Abstract

Acute kidney injury is common, with ∼13 million cases and 1.7 million deaths/year worldwide. A major cause is renal ischemia, typically following cardiac surgery, renal transplant or severe hemorrhage. We examined the cause of the sustained reduction in renal blood flow (“no-reflow”), which exacerbates kidney injury even after an initial cause of compromised blood supply is removed. After 60 min kidney ischemia and 30-60 min reperfusion, renal blood flow remained reduced, especially in the medulla, and kidney tubule damage was detected as Kim-1 expression. Constriction of the medullary descending vasa recta and cortical peritubular capillaries occurred near pericyte somata, and led to capillary blockages, yet glomerular arterioles and perfusion were unaffected, implying that the long-lasting decrease of renal blood flow contributing to kidney damage was generated by pericytes. Blocking Rho kinase to decrease pericyte contractility from the start of reperfusion increased the post-ischemic diameter of the descending vasa recta capillaries at pericytes, reduced the percentage of capillaries that remained blocked, increased medullary blood flow and reduced kidney injury. Thus, post-ischemic renal no-reflow, contributing to acute kidney injury, reflects pericytes constricting the descending vasa recta and peritubular capillaries. Pericytes are therefore an important therapeutic target for treating acute kidney injury.

## Introduction

The global burden of acute kidney injury is approximately 13 million cases a year (Ponce & Balbi, 2016). It is associated with a high mortality (1.7 million deaths per year, worldwide) (Gameiro et al., 2018; Hoste et al., 2018; Mehta et al., 2016), and COVID-19 has added to its incidence (Ronco et al., 2020). Renal ischemia followed by reperfusion, which can occur after cardiac surgery, renal transplant or severe hemorrhage, is the most common cause of acute kidney injury (Lameire et al., 2006; Lameire & Vanholder, 2001). Sustained renal blood flow reductions occur after ischemia and reperfusion, both in experimental studies and in patients after kidney transplantation (Cristol et al., 1996; Nijveldt et al., 2001; Ramaswamy et al., 2002). Following short periods of ischemia, blood flow to the renal cortex largely recovers following reperfusion, but medullary blood flow remains reduced for a prolonged period, especially in the hypoxia-sensitive outer medulla. Medullary no-reflow is a critical event for amplifying renal tissue injury following reperfusion (Conesa et al., 2001; Olof et al., 1991; Regner et al., 2009).

Renal no-reflow has been attributed to various causes, including impaired erythrocyte movement and leukocyte accumulation in renal capillaries, as well as increased intratubular pressure (Bonventre & Weinberg, 2003; Sutton et al., 2002; Wei et al., 2017; Yamamoto et al., 2002). However, after years of investigation, no effective treatment is available, even though no-reflow predicts a worse prognosis after kidney ischemia. We therefore investigated an alternative possible cause of no-reflow, i.e. ischemia-evoked contraction of pericytes that regulate capillary diameter, which might reduce renal blood flow and physically trap red blood cells. Indeed, in the brain and heart contractile pericytes on capillaries play a key role in reducing blood flow after ischemia (Hall et al., 2014; O’Farrell et al., 2017; Yemisci et al., 2009) because capillaries remain constricted by pericytes even when blood flow is restored to upstream arterioles. In the kidney, pericytes are associated with the cortical and medullary peritubular capillaries and the descending vasa recta. They play a key role in regulating renal medullary blood flow (Crawford et al., 2012; Pallone & Silldorff, 2001) which is a crucial variable for meeting the contradictory demands of preserving cortico-medullary osmotic gradients to allow water retention in the body, while maintaining adequate oxygen and nutrient delivery. This raises the question of whether pericytes also play a role in generating renal no-reflow after ischemia.

Few studies have investigated how ischemia affects renal pericytes (Kwon et al., 2008; McCurley et al., 2017; Zhang et al., 2018), and whether pericytes contribute to renal no-reflow. However, peritubular pericytes are damaged in cortical tissue of cadaveric renal allografts following ischemia-reperfusion (Kwon et al., 2008), suggesting that renal blood flow control may be disrupted after ischemia by pericyte dysfunction. Here we show that pericyte-mediated capillary constriction, especially of the descending vasa recta, makes a crucial contribution to no-reflow following renal ischemia and reperfusion. We further show that targeting pericyte-mediated constriction pharmacologically can reduce ischemia-evoked acute kidney injury.

## Results

### No-reflow after renal ischemia and reperfusion

We used a combination of laser Doppler perfusion measurements, low magnification imaging of blood volume, and high magnification imaging that resolved individual capillaries, to asssess the magnitude and cause of changes of renal perfusion after ischemia. Ischemia for 1 hour decreased perfusion of the renal medulla and cortex by ∼90% (both p<0.0001 vs. control; assessed with laser Doppler: Figure 1a ,b). After 30 min reperfusion, blood flow recovered to 49% of control (significantly reduced, *P* = 0.005, Figure 1a) in the medulla, but to 75% in the cortex (*P*=0.047, Figure 1b) (Regner et al., 2009). Perfusion was stable in the contralateral kidney throughout (Figure 1a, b). After 60 min reperfusion, medullary perfusion remained compromised at 40% of the control level (*P*=0.017, Figure S1), but cortical perfusion had fully recovered (to ∼20% above the control value, not significant, *P*=0.092, Figure S1). Despite this flow recovery, we show below that peritubular capillaries in the cortex can become blocked after ischemia.

**Figure 1:**
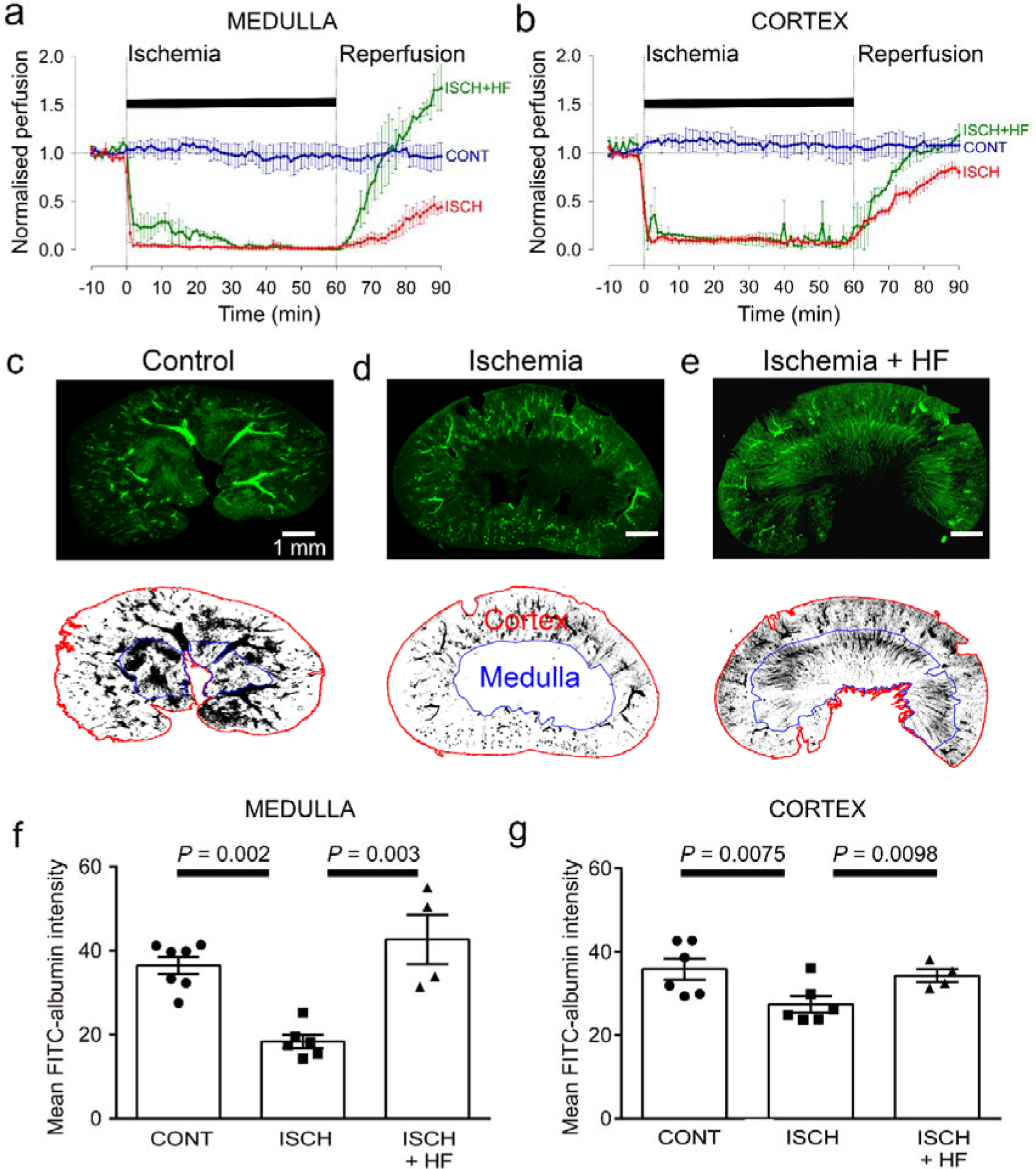
Ischemia and reperfusion lead to cortical and medullary no-reflow. **(a, b)** Ischemia (ISCH) evoked changes of blood flow (measured by laser Doppler) in the rat renal (**a**) medulla (n=4 animals) and (**b**) cortex (n=10 animals). CONT indicates blood flow on the contralateral (non-ischemic) side. Traces labeled +HF show the effect on recovery of perfusion of administering the Rho kinase inhibitor hydroxyfasudil (HF) immediately on reperfusion (ISCH+HF) (n=4 animals). **(c-e) Top:** low power views of kidney slices after perfusion *in vivo* with FITC-albumin gelatin, from (**c**) control (contralateral) kidney, (**d**) a kidney after ischemia and 30 min reperfusion, and (**e**) a kidney 30 mins after treatment with HF on reperfusion **Bottom:** regions of interest (ROIs) are shown in red and blue for the cortex and medulla. **(f)** Medullary perfusion (assessed in slices of fixed kidney as the total intensity of FITC-albumin summed over the ROIs) was reduced after 30 mins of post-ischemic reperfusion (51 stacks, 6 animals) by ∼50% compared with control kidneys (52 stacks, 7 animals). Treatment with HF increased medullary perfusion 2.3-fold at this time compared with non-treated ischemic kidneys (20 stacks, 4 animals). **(g)** Cortex perfusion (assessed as in c-e) after 30 mins of reperfusion after ischemia was reduced by ∼23.5% compared with control kidneys. Treatment with HF (ISCH+HF) increased cortex perfusion by 25% at this time compared with non-treated ischemic kidneys (ISCH). Data are mean±s.e.m. *P* values are corrected for multiple comparisons. Statistical tests used the number of animals as the N value.

After ischemia and reperfusion *in vivo*, assessing the volume of perfused vessels in fixed kidney slices, as the summed FITC-albumin intensity over ROIs, also demonstrated that renal ischemia and reperfusion led to no-reflow in the medulla compared with the non-ischemic kidney’s medulla (the perfusing blood volume was reduced by ∼50%, *P*=0.002; Figure 1c, d, f). Microscopic analysis resolving individual capillaries showed that this blood volume reduction was associated with a large reduction in capillary perfusion (Figure 2). The total perfused capillary length in 100 µm deep confocal z-stacks (frame size 640.17x 640.17 µm) was reduced by 35% (contralateral control 14689±3477 µm vs. ischemia 9527±1183 µm, *P*=0.038), the number of perfused capillary segments was reduced by 54% (control 530±82 vs. ischemia 244±30, *P*=0.03), and the overall perfused microvascular volume fraction was reduced by 51% (control 0.116±0.006 vs. ischemia 0.057±0.006, *P*=0.003; Figure 2e-g).

**Figure 2:**
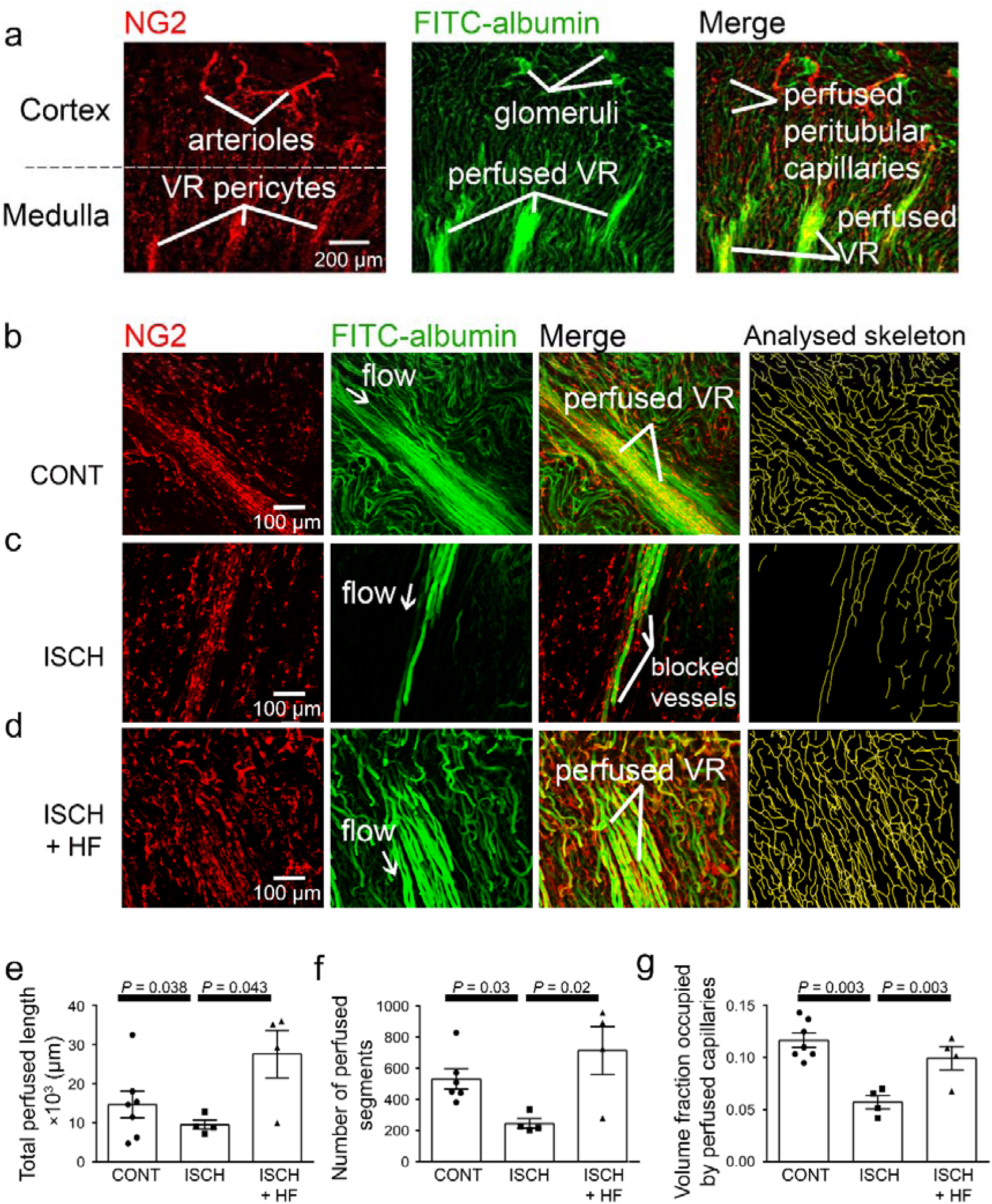
Ischemia and reperfusion reduce medullary microvascular perfusion. **(a)** Representative images of slices after perfusion with FITC-albumin gelatin, showing the rat kidney microcirculation in 100 µm deep confocal z-stacks. Images depict renal cortical arterioles, the glomeruli and peritubular capillaries, as well as the vasa recta capillaries (VR) that supply blood to the renal medulla. **(b-d)** Representative images of the medullary microcirculation: (**b**) in control conditions, (**c**) after ischemia and 30 mins reperfusion, and (**d**) after ischemia and reperfusion for 30 mins with hydroxyfasudil (HF) applied during reperfusion (ISCH+HF). Images show NG2-labelling of pericytes (red), FITC-albumin labelling (green) of vessels that are perfused, a merge of the NG2 and FITC-albumin images, and the analysed skeleton (yellow) of the perfused microvessels. (**e-g**) After ischemia and reperfusion (12 stacks, 4 animals), the total perfused capillary length **(e)**, the number of perfused capillary segments **(f)** and the overall volume fraction of vessels perfused **(g)** in 100 µm deep confocal z-stacks were reduced compared with control kidneys (14 stacks, 6-7 animals), and treatment with hydroxyfasudil immediately after reperfusion (10 stacks, 4 animals) increased all of these parameters. Data are mean±s.e.m. *P* values are corrected for multiple comparisons. Statistical tests used the number of animals as the N value.

In the cortex, perfusion was reduced less than in the medulla after ischemia and reperfusion, i.e. by 23.5% compared with non-ischemic kidneys (*P*=0.0075, Figure 1c, d, g). Furthermore, although a small percentage of afferent and efferent arterioles, and glomeruli, were not perfused in control conditions, this percentage did not increase significantly after ischemia (Figure 3a, b, g), and the arterioles’ diameter was not reduced compared with those in non-ischemic kidneys (Figure 3a, b, h, i). Similarly, it has been reported that upstream arteries are not constricted after ischemia (Yamamoto et al., 2002). In contrast, the total perfused peritubular capillary length in the 100 µm deep z-stacks (control 16441±1577 µm vs. ischemia 5411±2735 µm, reduced by 67%, *P*=0.03), the number of perfused capillary segments (control 550±32 µm vs. ischemia 349±54, reduced by 36.5%, *P*=0.01) and the overall perfused peritubular capillary volume fraction (control 0.12±0.01 vs. ischemia 0.06±0.02, reduced by 50%, *P*=0.01) were greatly reduced in the cortex when compared with non-ischemic kidneys (Figure 3d-f). Thus, the effect of ischemia and reperfusion is predominantly on the microvasculature, i.e. the peritubular cortical capillaries and the vasa recta, rather than on arteriolar segments of the kidney circulation. The Rho kinase data shown in Fig. 3 are discussed below.

**Figure 3:**
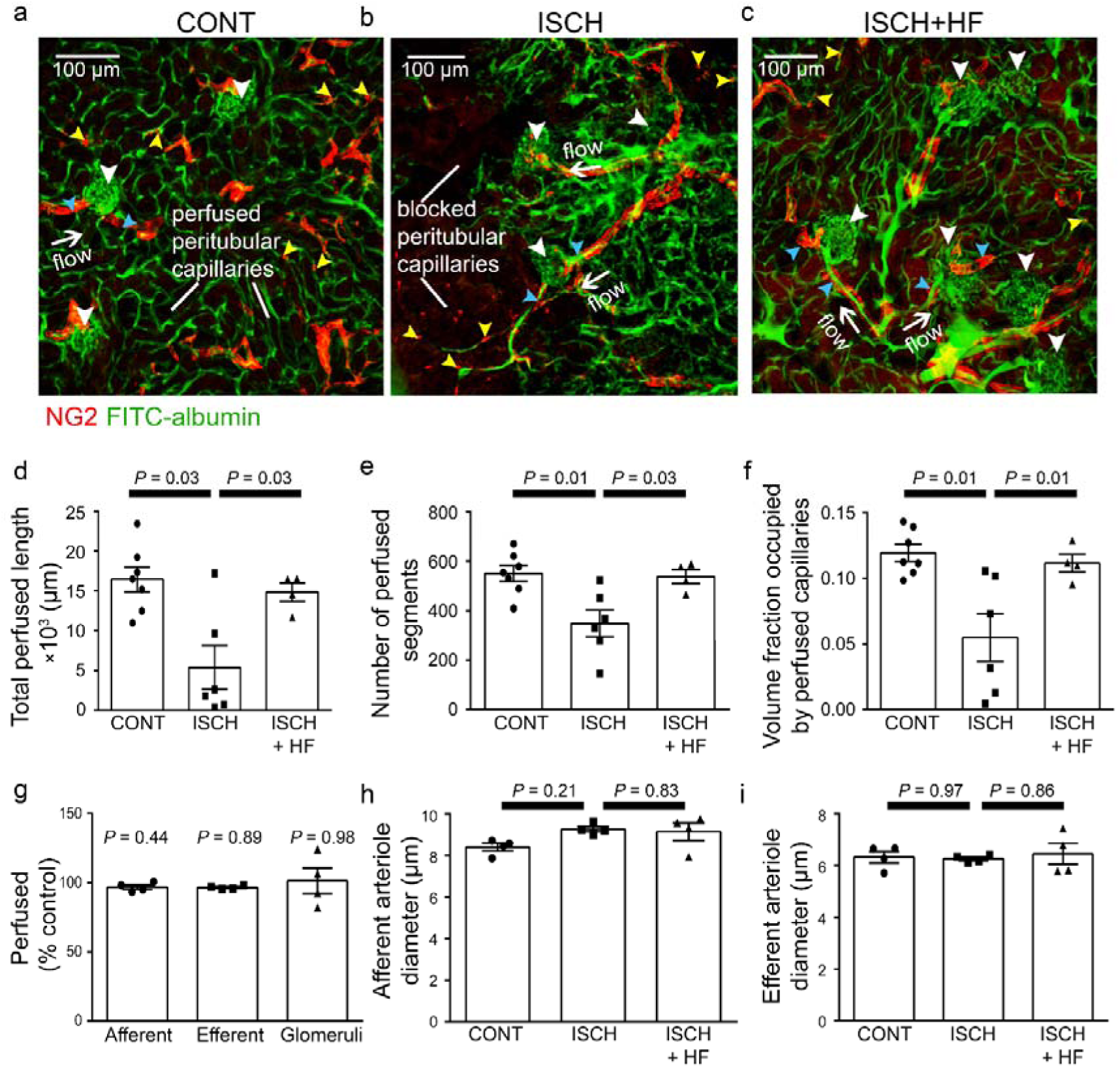
Ischemia and reperfusion of renal cortex evoke no-reflow in capillaries but not arterioles. (**a-c**) Representative images of rat renal cortex slices containing arterioles, glomeruli and peritubular capillaries, after perfusion with FITC-albumin gelatin: (**a**) for control kidneys (CONT), (**b**) after ischemia and reperfusion (ISCH), and (**c**) after ischemia with hydroxyfasudil (ISCH+HF). NG2-labelling is seen of arterioles and pericytes (red) (yellow arrowheads), while FITC-albumin labelling (green) shows vessels that are perfused. (**d-f)** After ischemia and reperfusion (12 stacks, 6 animals), the total perfused capillary length (**d**), the number of perfused segments (**e)**, and the overall perfused microvascular volume fraction (**f**) were reduced compared with control kidneys (14 stacks, 7 animals), and treatment with hydroxyfasudil immediately after reperfusion (10 stacks, 4 animals) increased cortical microvascular perfusion compared with non-treated ischemic kidneys. (**g**) Percentage of afferent and efferent arterioles (blue arrowheads in a-c), and of glomeruli (white arrowheads), perfused after ischemia, compared with control conditions. (**h-i**) Diameters of perfused (**h**) afferent and (**i**) efferent arterioles in the renal cortex for the three experimental conditions (15 arterioles, 4 animals for each group). Data are mean±s.e.m. *P* values are corrected for multiple comparisons. Statistical tests used the number of animals as the N value.

### Pericytes constrict descending vasa recta after ischemia and reperfusion

Higher magnification images demonstrated that, in control kidneys, only 9.7% of the descending vasa recta (DVR) capillaries were blocked (Figures 2b, 4d), i.e. were not perfused by FITC-albumin (Figures 2c, 4a-d). However, after ischemia and 30 mins reperfusion, 78% of the DVR capillaries were blocked (Figures 2c, 4a-d). Some capillaries were fully perfused and some completely unperfused throughout the area assessed, whereas some exhibited an abrupt cessation of blood flow with a decrease of FITC-albumin intensity over a few microns (Figures 2c, 4a-c). At block sites, the diameter of the FITC-albumin lumenal labeling at the final position blood reached was significantly lower in ischemic DVR capillaries compared with that at the much smaller number of block sites in non-ischemic controls (control 6.5±0.3 µm vs. ischemia 3.5±0.4 µm; *P*=0.039, Figure 4e). Thus, an ischemia-induced constriction of the DVR promotes blockage, which persists even after reperfusion.

**Figure 4:**
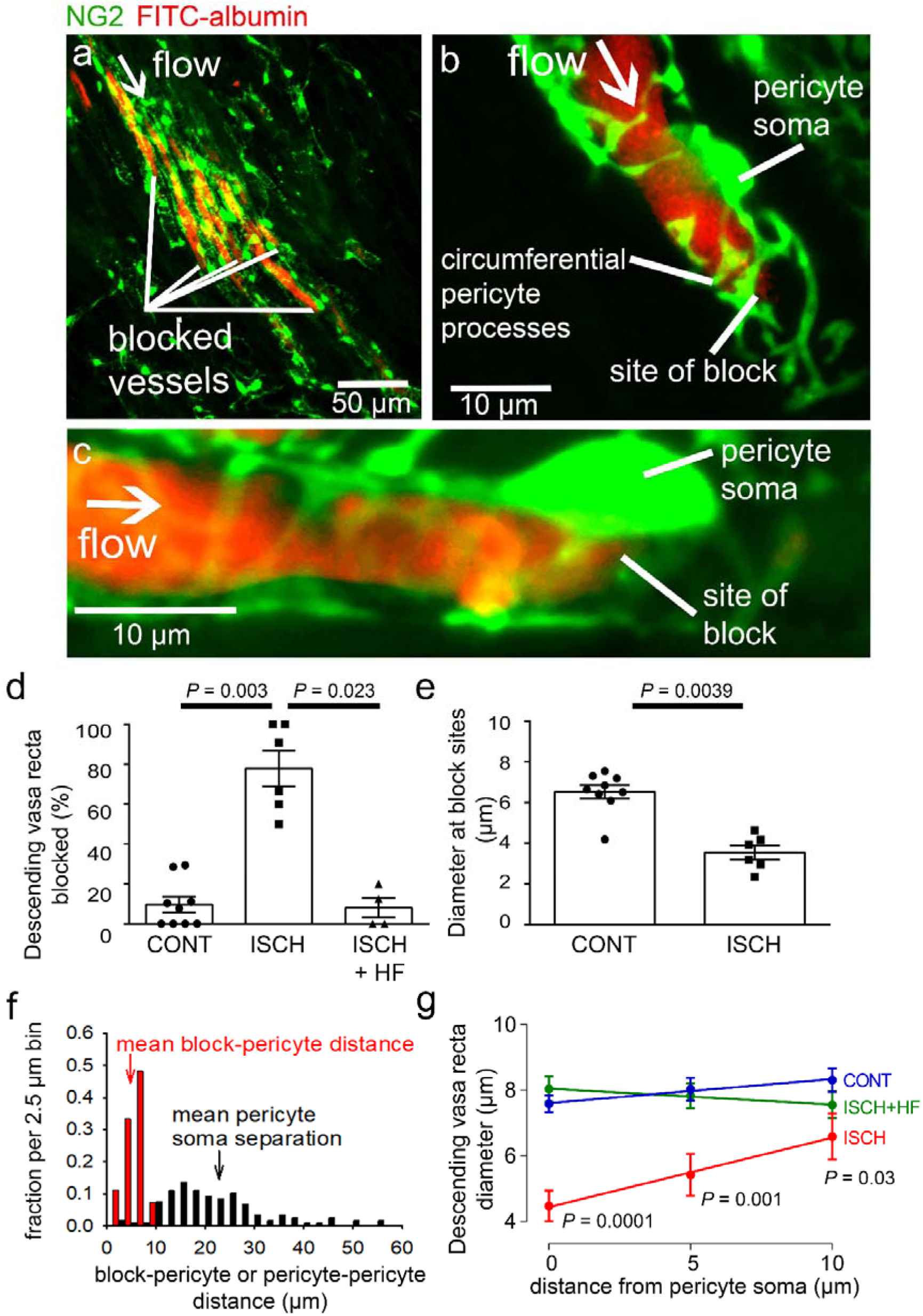
Descending vasa recta are constricted by pericytes after ischemia. **(a)** Descending vasa recta (DVR) in slices of rat renal medulla after perfusion with FITC-albumin gelatin (re-coloured red), and labelled for pericytes with antibody to the proteoglycan NG2 (green); FITC-albumin labeling shows perfused and blocked vessels. White arrow indicates flow direction; white lines indicate blocked vessels. **(b-c)** Representative images showing DVR capillaries blocked near pericyte somata. NG2-labelling of pericytes shows pericyte processes presumed to be constricting vessels at block site. **(d)** Percentage of DVR capillaries blocked in the renal medulla in control conditions (127 capillaries, 12 stacks, 9 animals), after ischemia and reperfusion (77 capillaries, 10 stacks, 6 animals), and after ischemia with hydroxyfasudil present in the reperfusion period (60 capillaries, 8 stacks, 4 animals). Statistical tests used number of animals as the N value. **(e)** Diameter at block sites. **(f)** Probability distribution per 2.5 μm bin of distance from blockage to nearest pericyte soma after ischemia and reperfusion (for 27 block sites), and of the distance between adjacent pericytes on DVR capillaries (for 118 pericyte pairs). **(g)** DVR diameter versus distance from pericyte somata (10 µm is approximately half the separation between pericytes) in the same 3 conditions as d (number of pericytes was 31, 20 and 17 respectively). *P* values by each point are from t-tests. Slope of the best-fit ISCH regression line is significantly greater than zero (*P*=0.039) while that of the CONT line is not (*P*=0.084). Data are mean±s.e.m.

Erythrocyte protein glycophorin A was labelled to assess if red blood cells were trapped at capillary regions of reduced diameter. Red blood cells were associated with only a small percentage of blockage sites in ischemic kidneys (5.8% of 85 blockages in 137 vessels from 2 animals), and even where red blood cells were near the capillary blockages they did not always block blood flow because FITC-albumin could pass the red blood cells (Figure S2a, b).

In the brain (Hall et al., 2014; Yemisci et al., 2009) and heart (O’Farrell et al., 2017) post-ischemic capillary constriction reflects pericyte contraction, which occurs near pericyte somata where circumferential processes originate (Nortley et al., 2019). From NG2 labelling we observed that many DVR blockages were close to pericyte somata, or near to pericyte circumferential processes connected to the soma (Figure 4b-c), suggesting that contraction of these juxta-somatic processes evoked capillary block. We measured the distance of 27 blockages to the nearest pericyte soma. The probability distribution of this distance is compared with that of the inter-pericyte distance in Figure 4f (if blocks did not depend on pericytes, the probability distribution of the blockage-pericyte distance would be constant until half the distance between pericytes). The mean blockage-pericyte distance was 4.87±0.33 µm after ischemia and reperfusion, which is less than a quarter of the distance between DVR pericytes (22.85±0.93 µm, from 118 pericyte pairs). Thus, these data are consistent with pericyte constriction generating the DVR blockages.

In control conditions, the few blockages occurring were mainly in regions where the inter-pericyte distance was larger. The mean distance from a blockage to the nearest pericyte soma was also larger (14.98±1.36 µm, p<0.0001 compared to post-ischemia), suggesting a different block mechanism in control conditions.

To assess pericyte-mediated DVR constriction further, we measured the FITC-albumin labelled lumen diameter at 5 micron intervals upstream of pericyte somata (upstream so there was FITC-albumin in the vessel: Figure 4g). After ischemia and reperfusion, the diameter was significantly reduced (by 41%, p=0.0001) near the pericyte somata compared with non-ischemic kidneys, but less reduced further from the somata. The diameter significantly increased with distance from the somata after ischemia and reperfusion (*P*=0.039 comparing the slope of the best-fit ischemia regression line with zero) but not in control conditions (*P*=0.084), implying constriction preferentially near the pericyte somata (Figure 4g) and identifying pericytes as the origin of the diameter reduction. Such constrictions will reduce blood flow directly by increasing the vascular resistance, and may also lead to blood cells becoming trapped at the regions of narrowed diameter, thus occluding the vessel and further reducing blood flow.

We assessed whether the endothelial glycocalyx (eGCX) contributed to DVR blockages. Labelling showed that eGCX is fairly uniformly present along capillaries, and this was not altered after ischemia (Figure S2f-g). There was no correlation between eGCX intensity and capillary diameter in control or ischemic conditions (Figure S2h). Thus, eGCX is not particularly associated with pericytes (Figure S2f), so the co-location of diameter reduction and blockages with pericyte somata presumably reflects pericyte process contraction rather than obstruction by eGCX.

### Pericytes constrict peritubular cortical capillaries *in vivo* after ischemia and reperfusion

Two-photon microscopy *in vivo*, of mice expressing dsRed in pericytes, revealed peritubular cortical pericytes constricting and blocking capillaries after ischemia and reperfusion (Figure 5a-c). This reduced the mean capillary diameter (averaged over all positions measured) from 10.8±0.2 to 8.1±0.5 µm (p<0.0001). To quantify whether ischemia-evoked blockages occurred disproportionately close to pericytes, we measured the distance of 15 blockages to the nearest pericyte soma. This distance was 4.12±0.39 µm, which is only 10% of the mean distance between peritubular cortical pericytes (41.3±2.6 µm, from 103 pericyte pairs). A plot of capillary diameter versus distance from pericyte somata (Figure 5d) showed that ischemia and reperfusion reduced the diameter by 40% at the somata (control 11.2±0.5 vs. ischemia 6.76±1.05 µm, *P*=0.001) with no significant effect on diameter far from the somata (control 10.3±0.2 µm vs. ischemia 9.6±0.5 µm, *P*=0.115). As in the medulla, the diameter increased significantly with distance from the pericyte somata after ischemia (*P*=0.046 comparing the slope of the best-fit regression line with zero) while in control conditions it did not (diameter decreased insignificantly with distance, *P*=0.10). Thus, capillaries are constricted specifically near cortical pericytes.

**Figure 5:**
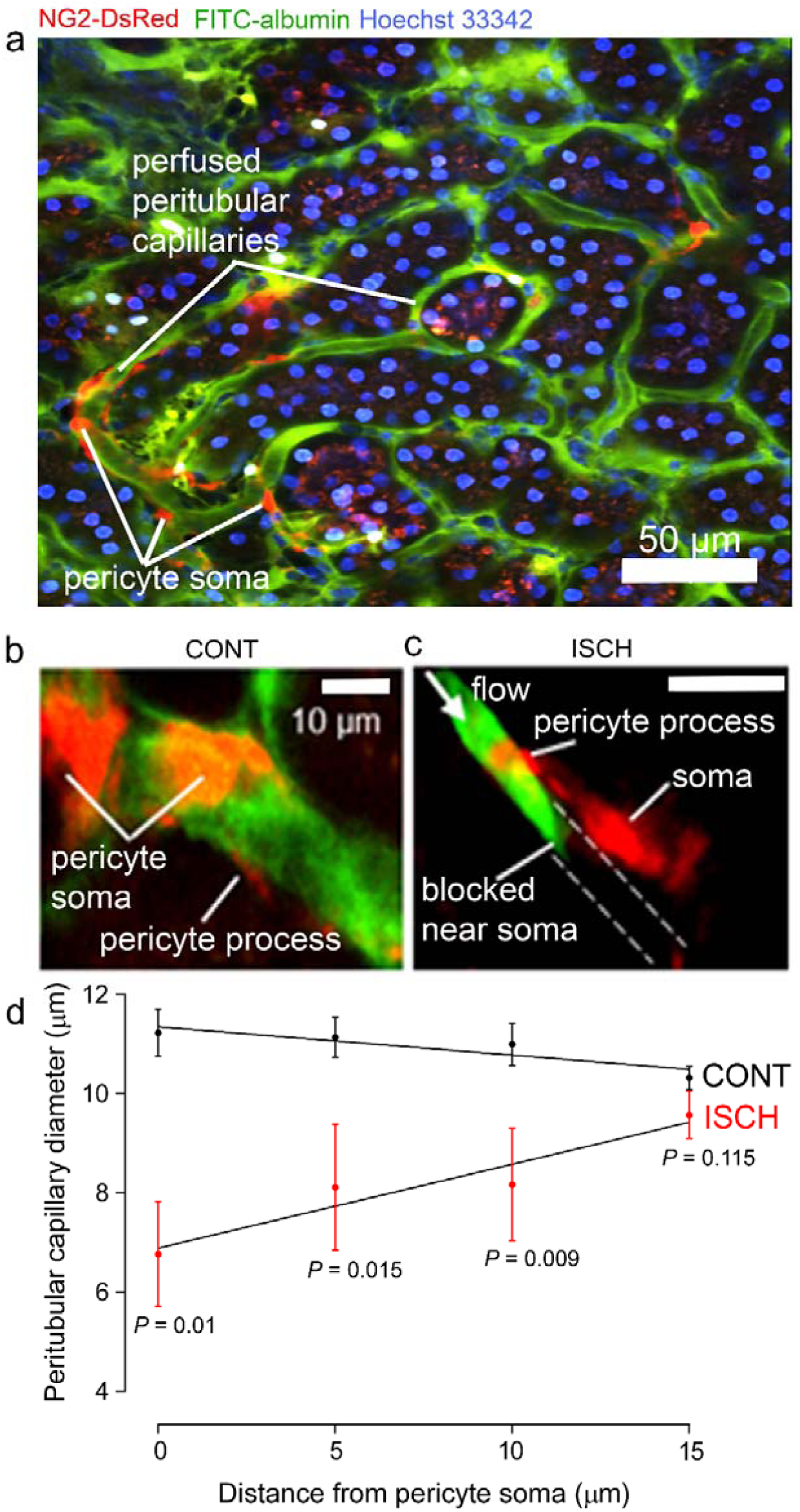
Pericytes constrict capillaries after renal ischemia *in vivo*. **(a)** Overview 2-photon *in vivo* imaging stack of the mouse renal cortex microcirculation, showing pericytes expressing NG2-DsRed (red), intraluminal FITC-albumin given intravenously (green), and Hoechst 33342 labelling nuclei (blue). Images were acquired in a plane parallel to the cortical surface. **(b, c)** Higher magnification images showing a pericyte on a cortical peritubular capillary in control conditions, and post-ischemic capillary block (dashed lines show path of blocked vessel). (**d)** Capillary diameter versus distance from pericyte somata after ischemia and reperfusion (ISCH), and for control kidneys (CONT) (number of pericytes was 15 and 10 respectively from 10 stacks from 3 animals from each group). Slope of the best-fit ISCH regression line is significantly greater than zero (*P*=0.046) while that of the CONT line is negative but not significantly different from zero (*P* = 0.10). Data are mean±s.e.m. *P* values comparing data at each distance are corrected for multiple comparisons. Statistical tests used number of images as the N value.

### Rho kinase inhibition reduces pericyte constriction and no-reflow

The contractility of pericytes depends partly on Rho kinase activity (Durham et al., 2014; Hirunpattarasilp et al., 2019; Homma et al., 2014; Kutcher et al., 2007). The Rho kinase inhibitor, hydroxyfasudil (3 mg/kg; i.v.), applied at the time of reperfusion to mimic a possible therapeutic intervention, significantly inhibited the decrease of renal medullary perfusion seen after ischemia-reperfusion (Figure 1a, e-f). *In vivo*, blood flow in the medulla (after 30 mins reperfusion) was increased 3.8-fold compared to ischemia without hydroxyfasudil (*P*=0.002, Figure 1a). Hydroxyfasudil induced a faster recovery of medullary blood flow than BQ123 (0.5 mg/kg, i.v.), an endothelin-A receptor antagonist (Figure S1c), but both resulted in blood flow at 30 mins reperfusion that was not significantly different from the control value (*P*=0.8 and 0.38 respectively) and was significantly higher than the flow seen after ischemia without either drug (*P*=0.01 for both drugs). In contrast, the angiotensin II type 1 (AT1) receptor antagonist valsartan (1 mg/kg i.v.) speeded the initial post-ischemic recovery of medullary blood flow, but did not return it to baseline by 30 mins reperfusion (Figure S1c). In the cortex, blood flow recovery on reperfusion was speeded by hydroxyfasudil and, after 30 mins of reperfusion, was increased 1.48-fold compared to ischemia alone (*P*= 0.02, Figure 1b). These data suggest that, in the medulla especially, activation of Rho kinase (in part downstream of ischemia-evoked activation of endothelin A receptors (Prakash et al., 2008; Wilhelm et al., 1999; Yamamoto et al., 2000)) contributes to ischemia-evoked pericyte-mediated capillary constriction.

Renal perfusion with post-ischemic inhibition of Rho kinase was also assessed in slices of fixed kidney (see above). Treatment with hydroxyfasudil during post-ischemic reperfusion prevented medullary no-reflow after ischemia and reperfusion: the blood volume was increased 2.3-fold compared to ischemia alone (*P*=0.003, Figure 1e-f), so that it did not differ significantly from that in control kidney (*P*=0.47). Hydroxyfasudil also increased ∼2.9-fold the total perfused medullary capillary length (*P* = 0.043), ∼2.9-fold the number of perfused capillary segments (*P*=0.02) and ∼2-fold the perfused volume fraction (*P*=0.0031) in medulla (Figure 2d-g). In the renal cortex, hydroxyfasudil given on reperfusion increased perfusion (blood volume) ∼1.25-fold (*P*=0.0098; Figure 1e, g), and increased the total perfused length of capillaries, the number of perfused capillary segments and the blood volume fraction to values that were not significantly different from those in non-ischemic kidneys (Figure 3c-f).

### Improvements of renal blood flow by hydroxyfasudil are via pericytes, not arterioles

Hydroxyfasudil might act on arteriolar smooth muscle or pericytes, or both. However, it had no effect on the diameter of afferent or efferent arterioles feeding and leaving the glomeruli (Figure 3h, i). In contrast, hydroxyfasudil reduced the constriction evoked at DVR pericyte somata by ischemia and reperfusion, increasing the diameter from 4.5±0.5 µm without hydroxyfasudil to 8.0±0.4 µm with the drug (p<0.0001) (Figure 4g), and reduced the percentage of DVR capillaries blocked from 78±9% to 8±5% (*P*=0.023), both of which are not significantly different from the values in non-ischemic kidneys (Figure 4d, f). Thus, ischemia induces, and hydroxyfasudil decreases, medullary no-reflow by specifically acting on DVR capillary pericytes rather than on upstream arterioles.

### Rho kinase inhibition reduces myosin light chain phosphorylation after ischemia

Rho kinase can inhibit (Riddick et al., 2008; Wang et al., 2009) myosin light chain phosphatase (MLCP), thus increasing phosphorylation of myosin light chain (MLC) by myosin light chain kinase (MLCK) and increasing pericyte contraction, but it also has other functions. To investigate how Rho kinase inhibition has the effects described above, we labelled for phosphorylated MLC. After ischemia and reperfusion, this was increased ∼11-fold for medullary and 5-fold for cortical pericytes (*P*=0.0001 in both locations, Figure 6a-j). Hydroxyfasudil treatment after reperfusion reduced this increase so that the labelling was not significantly different from that in control kidneys (*P*=0.95 and *P*=0.56 respectively; Figure 6a-j). Thus, if pericyte contraction is via conventional smooth muscle actomyosin, the reduced MLC phosphorylation could explain pericyte relaxation evoked by Rho kinase inhibition. Consistent with pericytes employing smooth muscle actomyosin, 56% of DVR pericytes near blockage sites labeled for the contractile protein α-SMA (Figure 6k-n; see also Park et al., 1997). Historically, demonstrating pericyte α-SMA labeling has been difficult, and a more favourable fixative might increase the percentage of cells labelled (Alarcon-Martinez et al., 2018).

**Figure 6:**
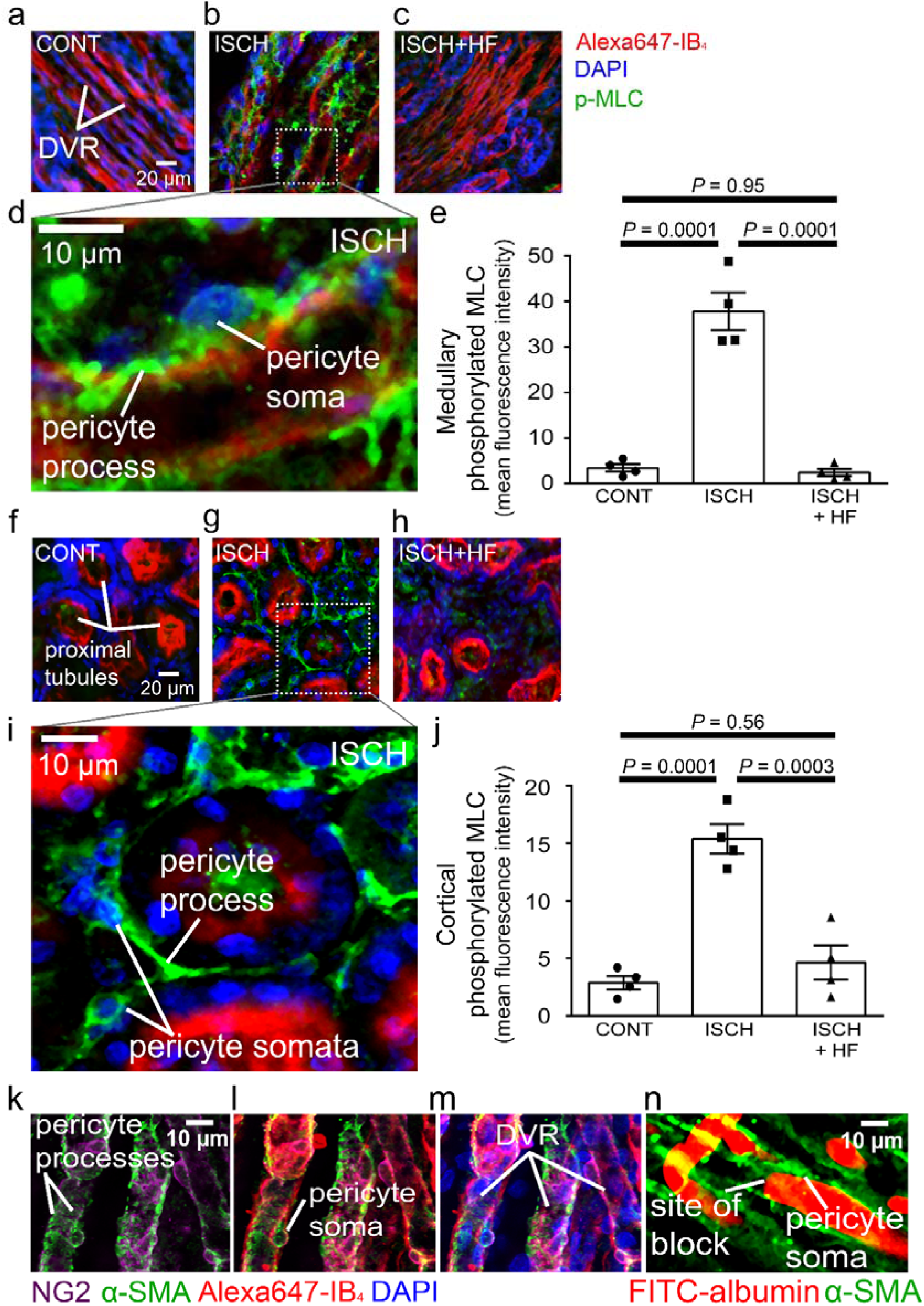
Pericyte contraction is mediated by _α_-SMA and regulated by Rho kinase. Representative images of the rat renal medulla containing descending vasa recta (DVR) pericytes **(a-d)** and cortical peritubular capillary pericytes **(f-i)**, labelled with antibody to phosphorylated myosin light chain (p-MLC, green), Alexa Fluor 647-isolectin B_4_ which labels kidney tubules and pericytes (red), and DAPI which labels nuclei (blue). Labelling is shown for kidneys in control conditions (CONT) **(a, f)**, after ischemia and reperfusion (ISCH) **(b, d, g, i)**, and after ischemia with hydroxyfasudil present during reperfusion (ISCH+HF) **(c, h)**. (**e, j**) Cortical (**e**) and medullary (**j**) p-MLC levels in pericytes for the three experimental conditions (10 stacks, 4 animals for each group, statistical tests used the numbers of animals for N values). **(k-m)** DVR pericytes labelled for NG2 (purple), α-SMA (green), Alexa647-isolectin B4 (red) and DAPI (blue). **(n)** DVR blockage-associated pericyte labelled for α-SMA. Data are mean±s.e.m. *P* values are corrected for multiple comparisons.

### Rho kinase inhibitor reduces reperfusion-induced acute kidney injury

Kidney injury molecule-1 (Kim-1) is a sensitive and early diagnostic indicator of renal injury in rodent kidney injury models (Vaidya et al., 2010), and in pathology is localized at high levels on the apical membrane of the proximal tubule where the tubule is most affected (Amin et al., 2004; Ichimura et al., 1998). Kim-1 levels in the proximal tubules were elevated 81-fold by ischemia and reperfusion (*P*=0.0004, Figure 7a, b, d), and treatment with hydroxyfasudil during reperfusion halved the Kim-1 labelling (*P*=0.03, Figure 7c, d).

**Figure 7:**
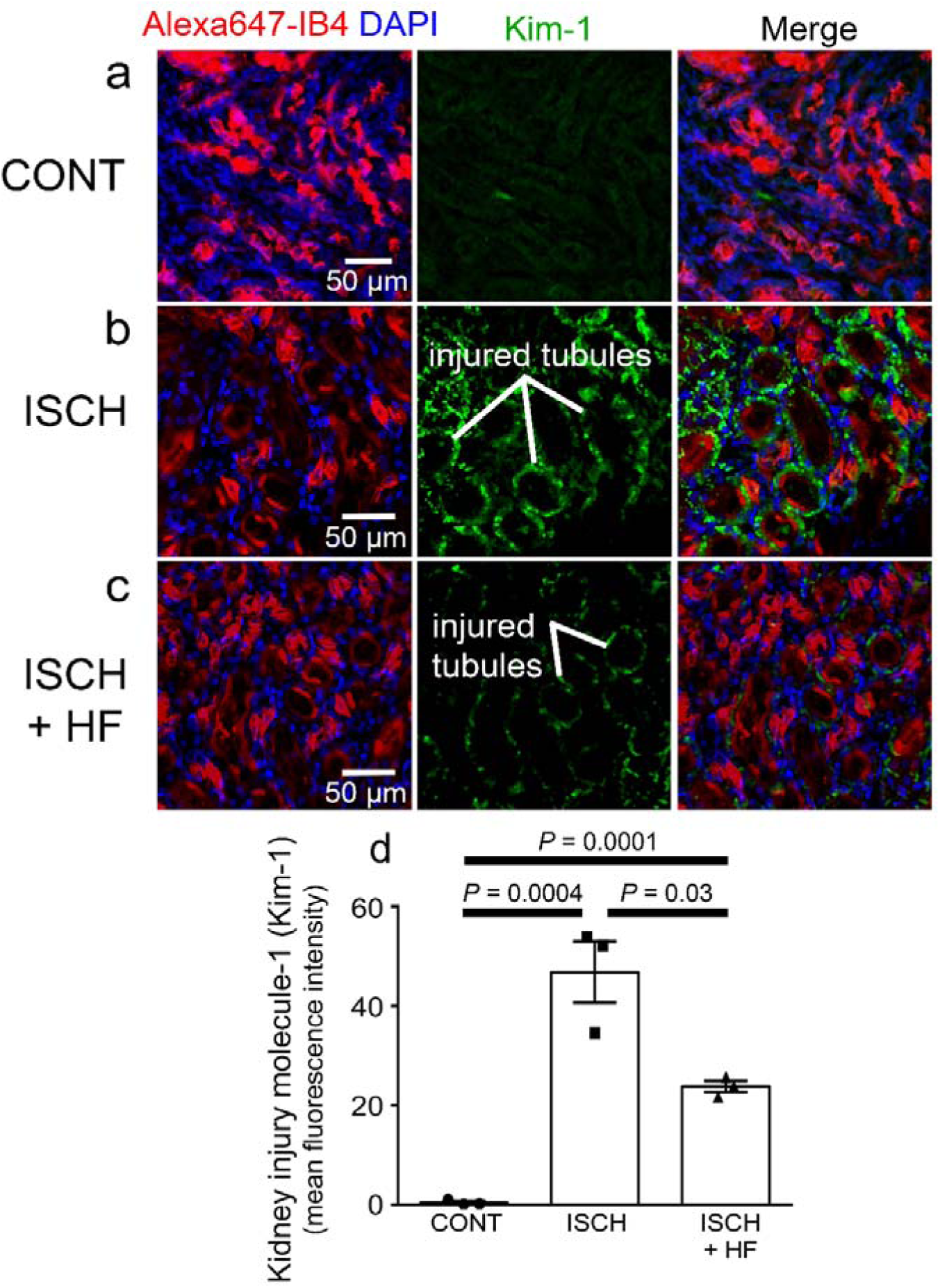
Rho kinase inhibition reduces kidney injury induced by ischemia and reperfusion. **(a-c**) Images of the rat renal cortex containing proximal tubules, showing isolectin B_4_ labelling kidney tubules (red), DAPI labelling nuclei (blue), and kidney injury molecule-1 (Kim-1) labelling as an injury marker (white lines indicate examples of injured tubules labelled in green), for control conditions (CONT) **(a)**, after ischemia and reperfusion (ISCH) **(b)**, and after ischemia with hydroxyfasudil present during reperfusion (ISCH+HF) **(c). (d)** Kim-1 levels for the three experimental conditions (6 stacks, 3 animals for each group). Data are mean±s.e.m. *P* values are corrected for multiple comparisons. Statistical tests used the number of animals as the N value.

## Discussion

This paper demonstrates, for the first time, that the long-lasting decrease of renal blood flow that follows transient ischemia is generated by pericyte-mediated constriction and block of the descending vasa recta and cortical peritubular capillaries, and that this post-ischemic no-reflow can be reduced pharmacologically. We found *in vivo* that sites of ischemia-evoked medullary and cortical capillary block were associated with pericyte locations. Furthermore, after ischemia and reperfusion, the diameters of descending vasa recta and peritubular capillaries were reduced specifically near pericyte somata, which extend contractile circumferential processes around the capillaries. In contrast, cortical arteriole diameters were not reduced and glomeruli remained perfused. The fact that capillary diameters are reduced specifically near pericyte somata establishes that this is due to a contraction of the circumferential processes of pericytes, and not (for example) due to a decrease in overall perfusion pressure (which would also reduce the diameter of capillaries away from pericyte somata). Together, these data establish pericyte-mediated capillary constriction as a major therapeutic target for treating post-ischemic renal no-reflow.

Pericyte-mediated constriction of renal capillaries may reflect reduced Ca^2+^ pumping in ischemia, raising [Ca^2+^]_i_ which activates contraction, as for CNS pericytes (Hall et al., 2014). Constriction may also partly reflect a release of angiotensin II (Allred et al., 2000; Boer et al., 1997; da Silveira et al., 2010; Miyata et al., 1999; Sanchez-Pozos et al., 2012; Zhang et al., 2004) and endothelin 1(Afyouni et al., 2015; Jones et al., 2020; Sanchez-Pozos et al., 2012) which raise [Ca^2+^]_i_ and Rho kinase activity (Lee et al., 2014; Shimokawa & Rashid, 2007), since we found that blocking endothelin 1 receptors and, to a lesser extent, angiotensin II receptors improved post-ischemic renal blood flow. Consistent with this, it has been demonstrated that vasoconstricting endothelin-A (Crawford et al., 2012; Wendel et al., 2006) and the angiotensin II type 1 (AT1) (Crawford et al., 2012; Miyata et al., 1999; Terada et al., 1993) receptors are located on pericytes along the descending vasa recta and regulate contractility at pericyte sites (Crawford et al., 2012). Additionally, endothelin 1 and angiotensin II evoke potent vasoconstriction of the descending vasa recta mainly through endothelin-A (Silldorff et al., 1995) and angiotensin II type 1 (AT1) (Rhinehart et al., 2003) receptors.

Rho kinase, a key downstream effector of both endothelin 1 and angiotensin II, inhibits the MLC dephosphorylation required to relax pericytes, thus promoting constriction (Hartmann et al., 2021). We found that blocking Rho kinase with hydroxyfasudil reversed ischemia-evoked pericyte-mediated capillary constriction, which could explain why Rho kinase block reduces acute kidney injury (Kentrup et al., 2011; Prakash et al., 2008; Teraishi et al., 2004; Versteilen et al., 2011; Versteilen et al., 2006), as we have confirmed using kidney injury molecule-1 (Kim-1) as a marker (Figure 7c, d). (In addition to inhibiting pericyte-mediated capillary constriction, hydroxyfasudil may also reduce kidney injury by reducing microvascular leukocyte accumulation, possibly by increasing the activity of endothelial nitric oxide synthase: Versteilen et al., 2011; Yamasowa et al., 2005). A direct effect of Rho kinase inhibition on pericyte contractility was confirmed by demonstrating that it reduced MLC phosphorylation in pericytes (Figure 6a-j) and that it increased capillary diameter specifically at pericyte somata (Figure 4g) in ischemic animals, implying that the effects of Rho kinase inhibition were on renal pericytes rather than an extra-renal systemic action. Hydroxyfasudil is the active metabolite of fasudil, a drug that has been clinically approved in Japan since 1995 for the treatment of vasospasm following subarachnoid hemorrhage (Lingor et al., 2019). Fasudil treatment improves stroke outcome in animal models (Vesterinen et al., 2013) and humans (Shibuya et al., 2005) and our data suggest that it may also be useful for reducing post-ischemic renal no-reflow and kidney damage.

We considered possible non-pericyte explanations for post-ischemic capillary constriction and block. Post-ischemic erythrocyte congestion in vasa recta has previously been described (Crislip et al., 2017; Olof et al., 1991) however red blood cells do not physically cause the capillary blockages observed after ischemia as they were associated with only a small percentage of block sites (Figure S2a, b). Thus, red blood cell trapping could be a consequence rather than a cause of the blockages. Leukocyte trapping may also contribute to reducing blood flow, but occurs on a longer time scale than we have studied (Kelly et al., 1994; Rabb et al., 1995; Ysebaert et al., 2000). Similarly, although a degradation of the eGCX has been reported after ischemia (Snoeijs et al., 2010; Song et al., 2018), we found a uniform distribution of the eGCX along the vessel wall, which was not modified after ischemia (Figure S2e-h), thus ruling out a causal association with capillary blockages which are preferentially located near pericytes. The present study demonstrates that pericyte-mediated constrictions of the descending vasa recta and cortical peritubular capillaries contribute to no-reflow and kidney injury at early stages of reperfusion, however we cannot exclude the possibility that other factors, such as inflammation and leukocyte infiltration (Gandolfo et al., 2009; Kelly et al., 1994; Rabb et al., 1995; Ysebaert et al., 2000), or eGCX dysfunction (Bongoni et al., 2019), might also contribute to post-ischemic microvascular injury at later phases of acute kidney injury. Furthermore, in response to the pericyte-mediated constriction evoked by ischemia, the DVR may undergo post-ischemic adaptations, releasing more nitric oxide at 48 hours post-ischemia which could reduce pericyte constriction at later times after ischemia than we have studied (Zhang et al., 2018).

The recovery of blood flow in the medulla on renal arterial reperfusion was slower than in the cortex. The regulation of renal medullary blood flow is mainly mediated by vasa recta pericytes, independent of total or cortical blood flow (Pallone & Silldorff, 2001). The need for accurate flow regulation in the relatively hypoxic medulla may account for pericytes on the DVR being much closer together (mean separation 22.9±0.9 µm) than for peritubular cortical pericytes (41.3±2.6 µm) and this may, in turn, contribute to a greater pericyte-mediated restriction of blood flow after ischemia in the DVR than in the cortical capillaries. Perhaps surprisingly, given our data, in post-cadaveric renal transplants a better outcome has been reported for kidneys with a higher number of pericytes immediately post-transplant (Kwon et al., 2008). This may, however, reflect an aspect of pericyte function other than capillary constriction, such as angiogenesis and maintenance of vessel integrity (Shaw et al., 2018), with these functions failing in transplanted tissue in which pericytes have already died due to ischemia.

Rodent models of renal ischemia can employ bilateral ischemia or unilateral ischemia with or without contralateral nephrectomy (Fu et al., 2018). In the present study, unilateral ischemia without contralateral nephrectomy (which may occur during renal-sparing surgeries) (Hollenbeck et al., 2006; Medina-Rico et al., 2018) was chosen to explore the early mechanisms of ischemia and reperfusion injury while using the contralateral kidney as a paired control for potential systemic hemodynamic changes that could be triggered during and after the surgical procedure. The presence of an uninjured contralateral kidney reduces animal mortality during the surgical procedure, and, thus, longer ischemia times can be used, resulting in more severe and reproducible injury (Fu et al., 2018; Le Clef et al., 2016; Polichnowski et al., 2020; Soranno et al., 2019). Unilateral ischemia-reperfusion without contralateral nephrectomy is considered a strong model to study the progression from acute renal injury to long-term tubulo-interstitial fibrosis (Fu et al., 2018; Le Clef et al., 2016; Polichnowski et al., 2020; Soranno et al., 2019), but we acknowledge that the model used in the present study may not be similar to some clinical situations where both kidneys are injured, and there are limitations of translatability from all animal models of acute kidney injury to human disease (Fu et al., 2018). A limitation of our study is that all experiments were performed on male rats and mice. Female rats are relatively protected against post-ischemic renal failure (Lima-Posada et al., 2017; Müller et al., 2002), possibly because in male rats androgens promote ischemic kidney damage by triggering endothelin-induced vascular constriction (Müller et al., 2002). However, these studies showed that sex did not influence ischemia repefusion-induced injury after 24 hours, but only after 7 days (Lima-Posada et al., 2017; Muller et al., 2002), i.e. on a much longer time scale than we have studied.

In the present study, we have shown that pericyte contraction contributes to reducing cortical and medullary blood flow at early stages of reperfusion. This initial pattern could also contribute to the pericyte injury, detachment and capillary rarefaction observed at later stages after ischemia and reperfusion (Kramann et al., 2017), which lead to further damage to the kidney (Khairoun et al., 2013; Kramann et al., 2017). However, there was no evidence of pericyte detachment during the time frame of the present study. Treatment from the beginning of reperfusion (to mimic a clinically-possible therapeutic approach) with hydroxyfasudil, a Rho kinase inhibitor, increased medullary and cortical blood flow, increased the post-ischemic diameter of DVR capillaries at pericyte locations, reduced the percentage of DVR capillaries that remained blocked, and reduced kidney injury after renal reperfusion. Presumably the protection of renal blood flow and downstream tissue health would be even greater if hydroxyfasudil could be given before ischemia was induced (e.g. in situations such as cardiac surgery and kidney transplantation, where renal ischemia might be anticipated). Thus, pericytes are a novel therapeutic target for reducing no-reflow after renal ischemia. Acute kidney injury caused by post-ischemic no-reflow causes significant socio-economic cost. Our identification of pericyte contraction as a therapeutic target for ischemia-induced acute kidney injury should contribute to the development or re-purposing of drugs that can prevent renal no-reflow.

## Methods

### Study approval

Experiments were performed in accordance with European Commission Directive 2010/63/EU and the UK Animals (Scientific Procedures) Act (1986), with approval from the UCL Animal Welfare and Ethical Review Body.

### Animal preparation for ischemia experiments

Due to the high density of kidney tissue, intravital microscopy is limited to superficial regions of the cortex <100 μm deep (Sandoval & Molitoris, 2017). As the renal medulla is inaccessible for in vivo imaging, we used laser Doppler flowmetry to assess blood flow changes of both kidneys or within the cortex and medulla of one kidney simultaneously. Additionally, we used FITC-albumin gelatin perfusion for measuring microvascular network perfusion (O’Farrell et al., 2017) in the renal cortex and medulla, supplemented with high resolution images of individual capillaries to assess the mechanisms underlying blood flow changes.

Adult male Sprague-Dawley rats (P40-50), or NG2-dsRed male mice (P100-120) expressing dsRed in pericytes to allow live pericyte imaging, were anesthetized with pentobarbital sodium (induction 60 mg/kg i.p.; maintenance 10-15 mg/kg/h i.v.). The femoral veins were cannulated to administer anesthetic and drugs. Stable kidney perfusion was confirmed using laser Doppler probes (OxyFlo™ Pro 2-channel laser Doppler, Oxford, United Kingdom) to measure blood flow in the contralateral kidney throughout the experiment, and anesthesia was monitored by the absence of a withdrawal response to a paw pinch. Body temperature was maintained at 37.0±0.5°C with a heating pad.

### Renal ischemia and reperfusion

Both kidneys were exposed, and the renal arteries and veins were dissected. Left kidneys were subjected to 60 min ischemia by renal artery and vein cross-clamp, followed by 30 or 60 min reperfusion. This reperfusion duration was chosen to assess pericyte function soon after starting reperfusion. Right kidneys underwent the same procedures without vessel clamping. Two laser Doppler single-fibre implantable probes of 0.5mm diameter (MSF100NX, Oxford Optronix, Oxford, United Kingdom) measured simultaneously the perfusion of both kidneys (or of the outer medulla and cortex of one kidney). Cortical and outer medullary perfusion were measured with the probe on or 2 mm below the kidney surface, respectively. Successful artery and vein occlusion was confirmed by a sudden fall of laser Doppler signal. Laser Doppler monitoring, which detects the movement of cells in the blood, is a widely used method for studies of microvascular perfusion in experimental and clinical studies and measures the total local microcirculatory blood perfusion in capillaries, arterioles, venules and shunting vessels (Fredriksson et al., 2009; Rajan et al., 2009). Laser Doppler is suitable for monitoring of relative renal microvascular blood flow changes in response to physiological and pharmacological stimuli in rodents (Lu et al., 1993; Vassileva et al., 2003).

Endothelial glycocalyx (eGCX) was labelled in vivo using wheat germ agglutinin (WGA) Alexa Fluor 647 conjugate (ThermoFisher, W32466, Waltham, MA) injected through the jugular vein (200 μl, 1 mg/ml) 45 minutes before renal ischemia/reperfusion (Kutuzov et al., 2018). WGA binds to N-acetyl-D-glucosamine and sialic acid residues of the eGCX. Using ImageJ, WGA fluorescence intensities were measured by drawing regions of interest (ROIs) across capillaries at the mid-points of pericyte somata, and away from the soma in 5 μm increments on both sides of the pericyte. Capillary diameters were also measured at each position.

Hydroxyfasudil hydrochloride, a reversible cell-permeable inhibitor of Rho kinase (Santa Cruz Biotechnology sc-202176, Dallas, TX) which is expected to decrease pericyte contractility (Hartmann et al., 2021; Kutcher et al., 2007) was administered as a bolus (3 mg/kg *i*.*v*.), immediately on starting reperfusion. This protocol, rather than having the drug present during the ischemic insult, better mimics a clinical situation where drugs could be given on reperfusion. Control and non-treated ischemic animals received saline infusion with the same volume.

### Animal perfusion and tissue preparation for imaging

After renal ischemia/reperfusion, animals were overdosed with pentobarbital sodium and transcardially-perfused with phosphate-buffered saline (PBS) (200 ml) followed by 4% paraformaldehyde (PFA, 200 ml) fixative and then 5% gelatin (20ml in PBS Sigma-Aldrich, G2625, Darmstadt, Germany) solution containing FITC-albumin (Sigma-Aldrich, A9771, Darmstadt, Germany), followed by immersion in ice for 30 minutes (adapted from (Blinder et al., 2013)). Kidneys were fixed overnight in 4% PFA, and 150 µm longitudinal sections made for immunohistochemistry. Rats have ∼64 ml of blood per kg bodyweight, thus the FITC-albumin gelatin solution would suffice to fill the total blood volume. The gelatin sets when the body temperature falls and traps FITC-albumin in the perfused vessels; blocked vessels show no penetration of FITC-albumin past the block.

### In vivo two-photon imaging

NG2-DsRed mice (P100-120) were anesthetized using urethane (1.55 g/kg i.p., in two doses 15 min apart). Anesthesia was confirmed by the absence of a paw pinch withdrawal response. Body temperature was maintained at 36.8±0.3°C. A custom-built plate, attached to the kidney using superglue and agarose created a sealed well filled with phosphate-buffered saline during imaging, when the plate was secured under the objective on a custom-built stage.

Peritubular capillary diameter was recorded during renal ischemia/reperfusion using two-photon microscopy of the intraluminal FITC-albumin (1 mg in 100 μl of saline given intravenously). Two-photon excitation used a Newport-Spectra Physics Mai Tai Ti:Sapphire Laser pulsing at 80 MHz, and a (Zeiss LSM710, Oberkochen, Germany) microscope with a 20× water immersion objective (NA 1.0). Fluorescence was excited using 920 nm wavelength for DsRed, and 820 nm for FITC-albumin and Hoechst 33342. Mean laser power under the objective was <35 mW. Images were analysed using ImageJ. Vessel diameter was defined using a line drawn across the vessel as the width of the intraluminal dye fluorescence.

### Immunohistochemistry

Pericytes were labelled by expression of DsRed under control of the NG2 promoter (in mice), or with antibodies to NG2 (1:200; Abcam ab50009, Cambridge, United Kingdom), α-smooth muscle actin (α-SMA) (1:100; Abcam ab5694, Cambridge, United Kingdom), or myosin light chain (phospho S20, 1:100, Abcam ab2480, Cambridge, United Kingdom), and the capillary basement membrane and pericytes were labelled with isolectin B_4_-Alexa Fluor 647 (1:200, overnight; Molecular Probes, I32450, Thermo Fisher Scientific, Waltham, MA). Z-stacks of the cortex and outer medulla (frame size 640.17×640.17 µm) for cell counting were acquired confocally (Zeiss LSM 700, Oberkochen, Germany). Pericyte intersoma distance was calculated between pairs of pericytes on capillaries within the same imaging plane. Kidney damage was assessed using kidney injury molecule-1 (Kim-1) antibody (1:100, overnight; Novus Biologicals, NBP1-76701, Abingdon, United Kingdom). Red blood cells were labelled with antibody to glycophorin A (1:2000, AbCam ab9520, Cambridge, United Kingdom). Alexa Fluor conjugated secondary antibodies were added overnight (1:500; ThermoFisher, A31572, A31556, A31570, Waltham, MA).

### Image analysis

Regions of interest (ROIs) were drawn around the renal cortex and medulla (Fig. 1), and the mean FITC-albumin signal intensity was measured for each ROI using ImageJ. This signal is assumed to provide an approximate measure of the amount of blood perfusing the tissue (conceivably downstream capillary constriction could lead to an upstream dilation and an increased blood volume being detected but, if this did occur, it would lead to an underestimate of the decrease of perfusion occurring). To gain a more accurate assessment of perfusion, we also used the ImageJ macro TubeAnalyst to measure the microvascular network of the renal cortex and medulla and obtain the total perfused capillary length, the number of perfused capillary segments and the overall perfused microvascular volume fraction. To quantify the percentage of perfused capillaries, we counted the number of filled (with FITC-albumin) and unfilled vessels that crossed a line drawn through the centre of each image perpendicular to the main capillary axis.

To assess whether pericytes cause flow blockages, we measured the distance along the capillary from the termination of the FITC-albumin signal to the mid-point of the nearest visible pericyte soma, since in brain most contractile circumferential pericyte processes (which can adjust capillary diameter) are near the pericyte soma (see Figures 4d, 5f, S2 and S3 of Ref (Nortley et al., 2019)). Capillary diameters were measured at the block sites where the FITC-albumin signal terminated. We also plotted the diameter of the FITC-albumin labelled capillary lumen as a function of the distance from the pericyte somata to assess whether diameter reduction was a nonspecific effect of ischemia, or was pericyte-related. A constriction seen specifically at pericyte somata is an unambiguous indication that pericyte contraction is occurring (Nortley et al., 2019). The identification, and direction of flow, of the afferent and efferent arterioles were deduced from tracking in confocal Z-stacks.

### Statistics

Statistical analysis employed Graphpad Prism (San Diego, CA). Data normality was tested with Shapiro-Wilk tests. Normally distributed data were compared using Student’s 2-tailed t-tests or ANOVA tests. Data that were not normally distributed were analysed with Mann-Whitney or Kruskal-Wallis tests. *P* values were corrected for multiple comparisons using a procedure equivalent to the Holm-Bonferroni method or Dunn’s test (corrected *P* values are significant if they are less than 0.05).

## Author contributions

FF devised experiments, carried them out, analysed data and wrote the first draft of the paper. DA helped to devise experiments and analyse data, and edited the paper.

## Acknowledgments

We thank Jonathan Lezmy, Svetlana Mastitskaya and Thomas Pfeiffer for comments on the manuscript.

---

**Figure S1:**
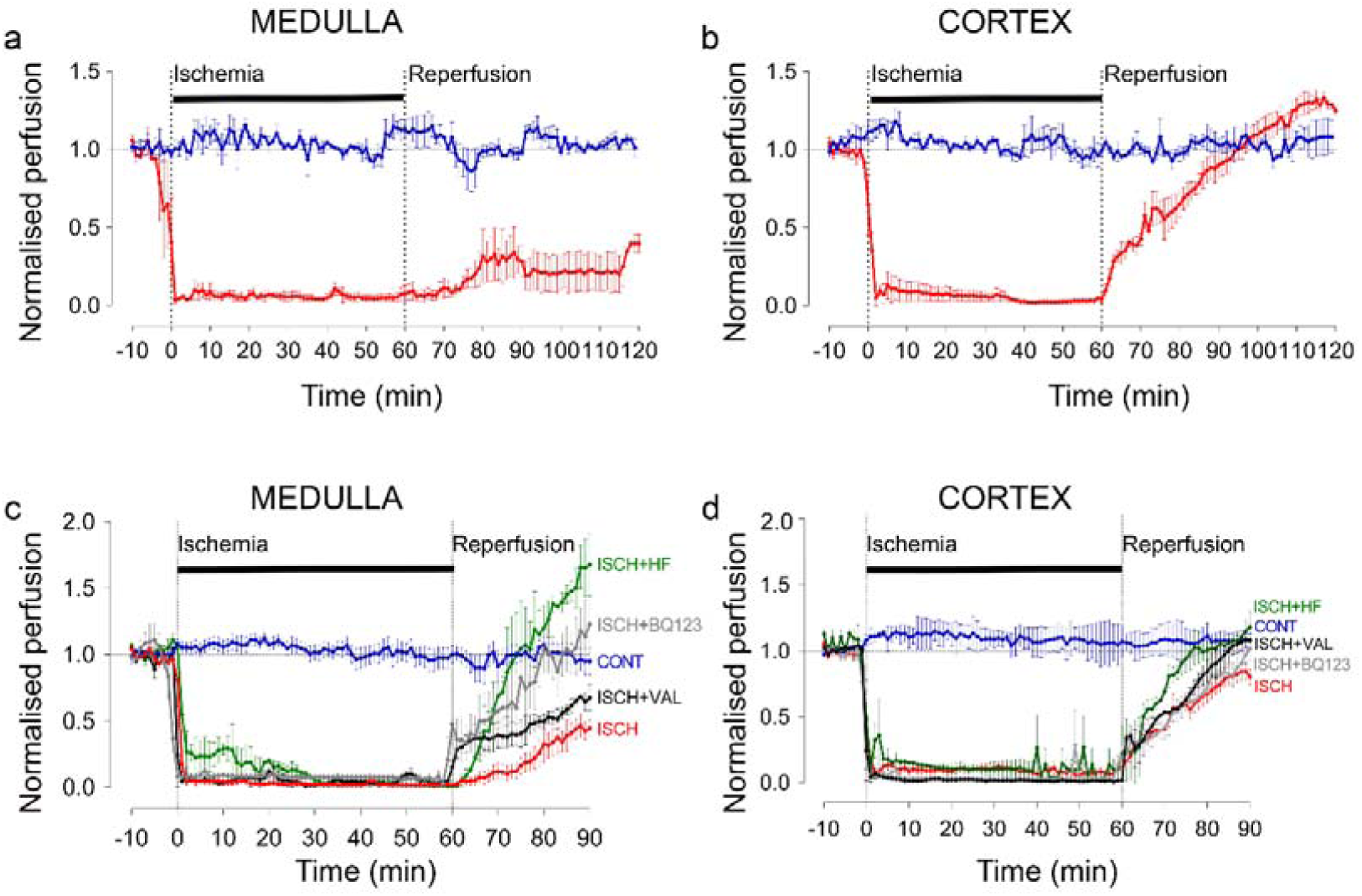
**(a, b)** Ischemia (ISCH) evoked changes of blood flow (measured by laser Doppler) in the rat renal (**a**) medulla (n=3 animals) and (**b**) cortex (n=3 animals). CONT indicates blood flow on the contralateral (non-ischemic) side. At 60 min following reperfusion, medullary perfusion remained compromised at 40% of its control value (*P*=0.017), but cortical perfusion was fully recovered (to ∼20% above the control value, although this did not reach significance, *P*=0.092). (c) Hydroxyfasudil (3 mg/kg; i.v.) treatment immediately after reperfusion (ISCH+HF) induced a faster recovery to the pre-ischemic value of of medullary blood flow than did BQ123 (0.5 mg/kg, i.v., given on reperfusion: ISCH+BQ123), a selective endothelin-A receptor antagonist. After 30 min reperfusion both agents resulted in blood flow that was not significantly different from control (*P*=0.8 and *P*=0.38, respectively) but was significantly different from ischemia (*P*=0.01 for both drugs). Valsartan (1 mg/kg i.v., given on reperfusion: ISCH+VAL), an angiotensin II type 1 (AT1) receptor antagonist, increased medullary perfusion by 52% after 30 mins reperfusion compared with non-treated ischemic kidneys, although this did not reach significance (*P*=0.11 vs. ISCH) and valsartan had not reversed medullary blood flow to the baseline level after 30 mins (*P*= 0.19 vs. CONT). (d) Recovery of cortical blood flow to its control level on reperfusion was faster in the presence of hydroxyfasudil (ISCH+HF). BQ123 (*P*=0.05 vs. ISCH) and valsartan (*P*=0.04 vs. ISCH) also promoted recovery of cortical blood flow at 30 min reperfusion compared with non-treated ischemic kidneys (ISCH). Statistical tests used the number of animals as the N value.

**Figure S2:**
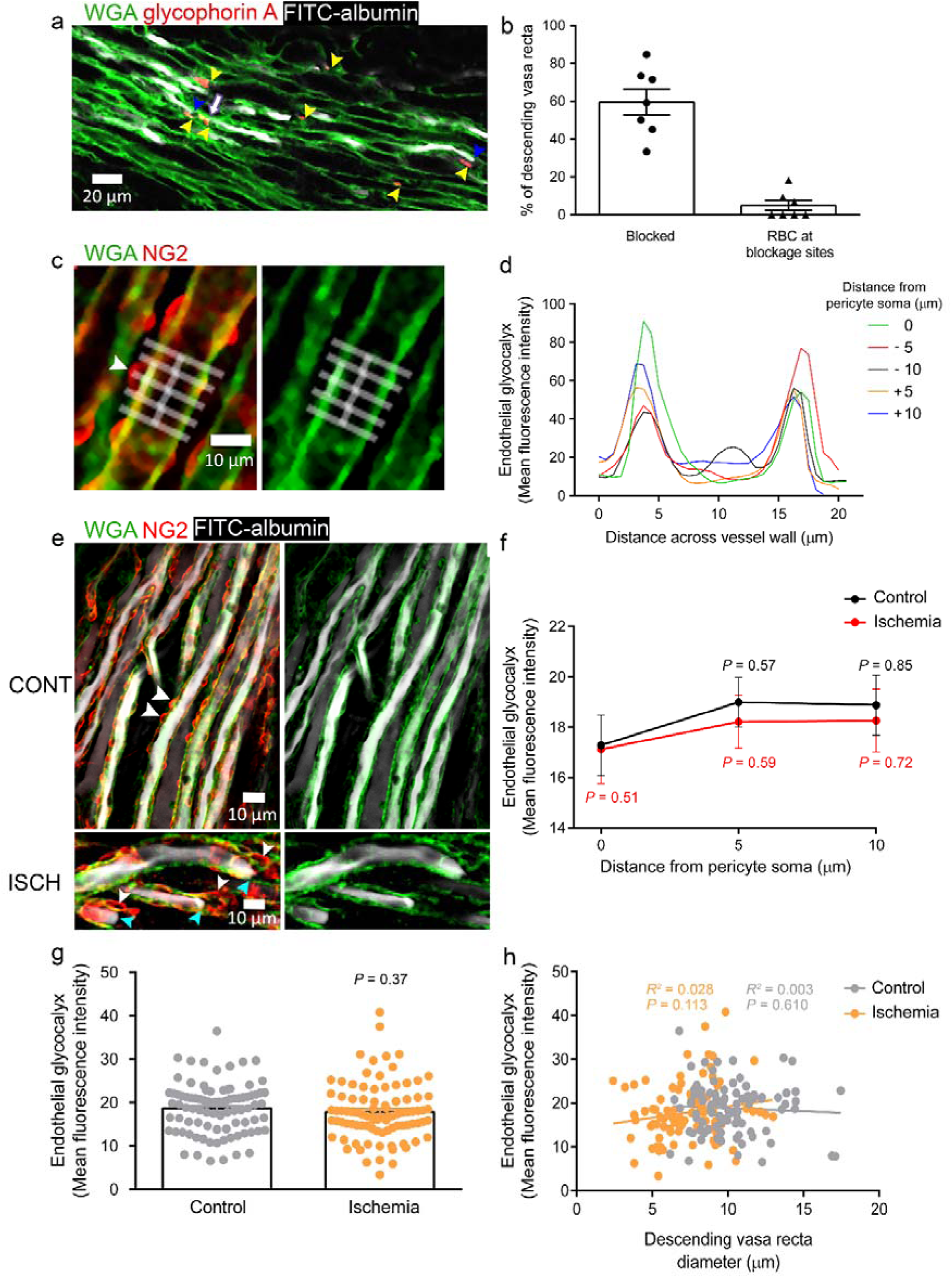
(**a**) Red blood cells (RBCs, indicated by yellow arrowheads, labelled for glycophorin A) were associated with a small percentage of blockage sites (indicated by blue arrowheads) in ischemic rat kidneys (5.8% of 85 blockages from 137 vessels analysed from 2 animals), and even where red blood cells were near the capillary blockages it did not always lead to a block of blood flow (as shown by FITC-albumin, re-coloured white, passing the red blood cells [purple arrow]). Note that the vasculature was perfused with PBS to remove loose RBCs before perfusing PFA and FITC-albumin, so the only RBCs remaining should be those bound to the vessel walls. (**b**) Percentage of DVR that were blocked, and percentage of blocked DVR that had an associated RBC. (**c**) Endothelial glycocalyx (eGCX) was labelled in vivo using wheat germ agglutinin-Alexa Fluor 647 (WGA, re-coloured green). White boxes show ROIs for measuring eGCX mean fluorescence intensities at different distances from the pericyte soma. (**d**) Plots of WGA signal across capillary at different distances from arrowed pericyte in (c). (**e**) eGCX is fairly evenly distributed along the vessel wall in normal kidneys, and also after ischemia and reperfusion. Blockages (indicated by blue arrowheads) are highly associated with pericyte location (indicated by white arrowheads) in ischemic kidneys (ISCH). (**f**) Mean level of eGCX averaged across vessel at different distances from the pericyte soma in control kidney and after ischemia with 30 mins reperfusion. For the control condition, black *P* values compare the value at each position with that at the soma. Red *P* values compare the ischemic and control groups for each position). (**g**) eGCX mean fluorescence averaged over all positions measured. (h) eGCX intensity and diameter have no correlation in control or ischemic conditions. Data are mean±s.e.m, 30 pericytes from 2 animals for each experimental condition. Statistical tests used the number of pericytes as the N value.

